# Development of a Novel Single Cell Multiomics Approach for Simultaneous Analysis of Replication Timing and Gene Expression

**DOI:** 10.1101/2025.01.02.631160

**Authors:** Anala V. Shetty, Clifford J. Steer, Walter C. Low

## Abstract

Replication timing (RT) allows us to analyze temporal patterns of genome-wide replication, i.e., if genes replicate early or late during the S-phase of the cell cycle. RT has been linked to gene expression in normal and diseased acute and chronic states such as cancer. However, studies done to date focused on bulk cell populations that required tens of thousands of cells for RT analysis. Here, we developed an affordable novel single cell (sc)-multiomics approach to simultaneously analyze RT and gene expression from cells or nuclei. We used this approach to generate sc-RT profiles and sc-gene expression data from the well-established human liver cancer cell line, HepG2. We demonstrated that as few as 17 mid S-phase cells were sufficient to produce cell-type specific pseudo bulk RT profiles that had a high correlation to previously published HepG2 bulk RT profiles. The sc-RT profiles allowed us to visualize how individual cells progressed through genome replication. We were also able to demonstrate high-resolution correlations between RT and gene expression within each individual cell, which to our knowledge, has not been reported. We observed trends that were conserved between individual cells, as well as cell-to-cell variations, which were not possible to detect with the bulk RT studies.

## Introduction

Replication timing (RT) of cells provides insight into how different parts of the genome replicate during S-phase of the cell cycle. We can deduce if individual genes replicate during the early, mid, or late S phase of the cell cycle. It is conserved within cell types, mitotically inherited (Hiratani & Gilbert, 2009), and is regulated in mega base sized bins throughout the genome (Pope et al., 2010). RT of the gDNA is disrupted in diseased states such as premature aging (Rivera-Mulia et al., 2017), cancer (Liu et al., 2013) and other acute and chronic conditions. Multiple cancer studies have shown that alterations in RT vary from small location-specific RT changes to larger chromosome wide delays in RT, as observed in 80% of cancers over the last century (Donley & Thayer, 2013; Vouzas & Gilbert, 2023). Although RT is affected in cancer cells, the cause or consequence for altered RT is not clear (Briu et al., 2021).

RT has also been correlated with gene expression in both normal and diseased states. Studies have demonstrated that early replicating genes are usually upregulated, and late replicating genes are downregulated and/or not expressed (Hiratani et al., 2008; Rivera-Mulia et al., 2015). The ability to study RT and its associations with gene expression, at the level of individual cells, would be powerful in understanding how these multilayer controls exist in normal and diseased cells.

We developed a novel single cell multi-omics approach that enables simultaneous analysis of RT and the accompanying transcriptome, from the same single cells, which to our knowledge. This allows us to observe cell-to-cell variation in RT and gene expression, as well as perform RT-gene expression correlations within individual cells. For this study, we chose a well-established human HepG2 cancer cell line (Knowles et al., 1980). HepG2 is a model cell line for studying human hepatocellular carcinoma cancer as well as primary hepatocytes *in* vitro (Štancl et al., 2024). These cells are easy to grow, and our lab had previously generated bulk RT data for HepG2 cells (Meyer-Nava et al., 2023). In processing HepG2 cells using the single cell multi-omics protocol, we analyzed (i) genome-wide coverage and copy number variations (CNVs); (ii) single cell RT and gene expression; and (iii) correlation between RT and gene expression using pseudo bulk profiles and within individual cells. For the first time, we demonstrated RT-gene expression correlations within the same cells, and at the single cell level. The results of our study emphasize the importance of the sc-multiomics approach in capturing high resolution relationships and associations between the gDNA and mRNA.

### Development of the single cell multiomics protocol

Protocols for generating either sc-RT or sc-RNA profiles exist, but simultaneous analysis of both parameters, from the same cells, has not been reported.

Advantages of our in-house sc-multiomics protocol include, (i) compatibility with both cells and nuclei, (ii) compatibility with very small cell numbers (as low as 10-20 cells) that can be used to process both genome and transcriptome per cell from scarce patient samples/limited cell numbers; and (iii) affordability vs. commercially available kits (∼15% of the cost of commercial kits for processing sc-gDNA and scRNA).

We were inspired by protocols like the G&T protocol (Macaulay et al., 2015) and SMARTSeq2 technique (Picelli et al., 2014) and adopted/combined their strategies for separating gDNA and mRNA from each cell, and for processing mRNA. For amplification of gDNA, we developed an affordable in-house gDNA protocol which can also be used as a stand-alone protocol. This gDNA protocol is more compatible with the newer 2-color chemistry Illumina sequencing platforms such a NovaSeq and NextSeq, compared to previously published in-house gDNA protocol (Bartlett et al., 2022); and it significantly reduces sequencing depth per sample for generating sc-RT profiles. This protocol can be used for detecting sc-RT and sc-CNV at the level of individual genes within both normal and diseased cells, such as those in cancer. The sc-gDNA processed using this protocol was compatible with two commonly used computational sc-RT pipelines, the Kronos sc-RT pipeline (Gnan et al., 2022) and the sc-Repliseq pipeline (Miura et al., 2020).

Bulk RT protocols generally use 20,000 to 40,000 cells per replicate of the G1, Early S, Late S, and G2 populations (with 3 replicates per experiment) (Marchal et al., 2018). With the in-house protocol, we demonstrated that as few as 17 single HepG2 mid S-phase nuclei/cells were sufficient to deduce pseudo bulk RT profiles that demonstrated high correlation with the bulk RT profile. This demonstrated the unique ability of the current protocol to drastically reduce cell numbers, while providing high resolution cell-specific and gene-specific information.

### Separation of gDNA and mRNA from single cells

The detailed steps for separation of gDNA from mRNA within each cell are described in the methods. The separation of gDNA from mRNA was adopted from the G&T protocol with minor modifications (Macaulay et al., 2016). Briefly, single cells/nuclei were sorted into RLT buffer in 96-well plates, with 1 cell per well. The single cells disintegrated in the RLT buffer releasing the gDNA and the RNA into the solution. The mRNA in each well was pulled down using the SMART-seq2 technique of using poly-dT tailed primers attached to magnetic beads (Picelli et al., 2014). The poly-dT primers were complementary to the poly-A tails of the mRNA. The beads were pulled down using a low elution magnet, leaving the gDNA in the supernatant (**Fig. 1**). The supernatant containing the gDNA was then separated into a new 96-well plate, maintaining the cell identity per well.

**Figure 1:**
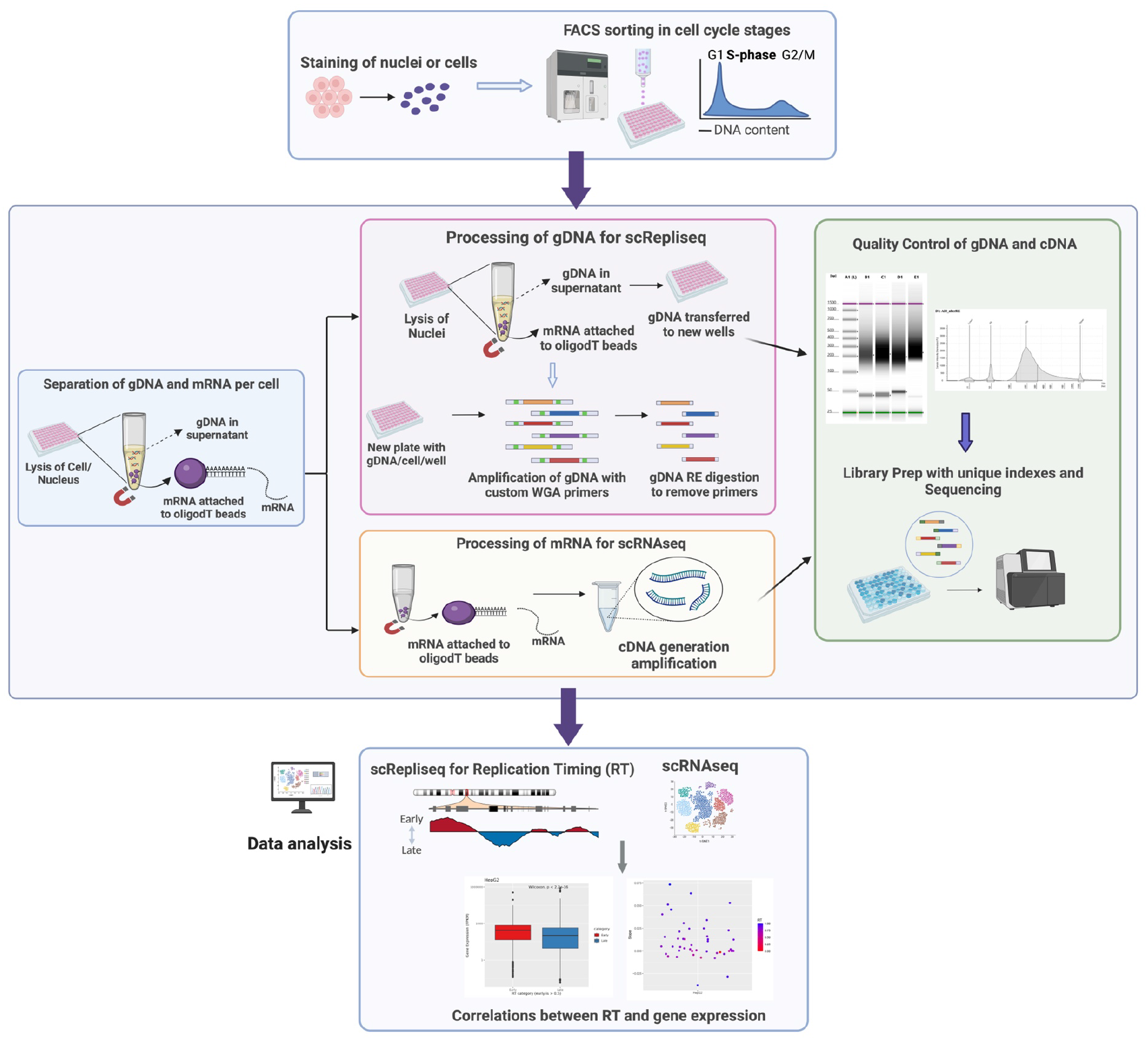
Single Cell Multiomics Approach for Determining Replication Timing and Gene Expression. Single cells (or nuclei) were stained with a DNA dye and then sorted into 96-well plates, based on their cell cycle. From each individual cell, both gDNA and mRNA were isolated, separated, and amplified using in-house protocols. Quality control of amplified gDNA and cDNA (derived from mRNA) was performed to ensure successful amplification, after which the samples were library prepped for sequencing. After sequencing, data analysis was performed to obtain information regarding copy number variation, replication timing (RT), and gene expression from each cell. High resolution correlations between RT and gene expression were drawn within each cell.

### Amplification of single cell gDNA using an *in-house* protocol

We developed an in-house protocol for processing sc-gDNA from either cells or nuclei. This protocol can be used for processing only gDNA from samples or can be used as part of the multiomics protocol. If performing the multiomics approach, we first pulled down the gDNA from the gDNA supernatant onto AMPure XP beads (**Supplemental Fig. 1A**). When processing only gDNA, cells/nuclei were directly sorted into the L&F buffer and processed. In both cases (with or without beads), we followed the same subsequent processing steps as described in the methods. We used a degenerate oligonucleotide primed (DOP)-PCR technique for amplification. We designed WGA oligos that contained RE digestion sites specific to the enzyme AcuI. After gDNA amplification, we performed RE digestion to remove the majority of primer sequences from both ends of the amplified DNA (**Supplemental Fig. 1A**). This drastically reduced presence of the high G% and repeat regions in the final gDNA fragments of interest. This protocol is more compatible with the newer Illumina sequencing machines that use two-color chemistry (eg. NextSeq, NovaSeq), compared to the Bartlett protocol (Bartlett et al., 2022) which was only compatible with the four-color chemistry sequencing platforms (eg. HiSeq (discontinued), MiSeq).

The two-color sequencing machines have filters that automatically discard low diversity sequences, such as the high % ‘G’ repeat regions on the WGA primers. Sequencing of the gDNA fragments without RE digestion led to the machine automatically discarding the majority of reads, due to the low diversity primers in the first 25 cycles. This issue was observed when sequencing gDNA processed using the Bartlett protocol (Bartlett et al., 2022), which was only compatible with the 4-color chemistry sequencing machines. This was addressed and resolved by using the RE digestion step in the protocol.

### Amplification of single cell mRNA

After the separation step using poly dT tailed magnetic beads, the mRNA was first converted into cDNA as described in the methods. Reverse transcription and amplification of mRNA was performed in-house using the strategy in G&T with minor modifications (Macaulay et al., 2016). We performed quality control to ensure that the amplified cDNA was in the size range expected before tagmentation-based library preparation as described.

### Processing HepG2 nuclei using the single cell multiomics protocol

#### Sorting HepG2 nuclei based on cell cycle phase

For the multiomics protocol, we processed HepG2 nuclei. We isolated nuclei from cells and sorted them based on DNA content using a previously optimized in-house protocol (Meyer-Nava et al., 2023). We stained the isolated nuclei with Vybrant DyeCycle Violet which is a DNA-specific dye (**Fig. 2A**). This allowed us to sort cells from specific phases of the cell cycle, based on DNA content (**Fig. 2C**). Cells in the G1 phase have 2 copies of chromosomes, cells in G2 have 4 copies, and the population in between is in the S phase, where the cells are actively replicating (>2 copies but <4 copies).

**Figure 2:**
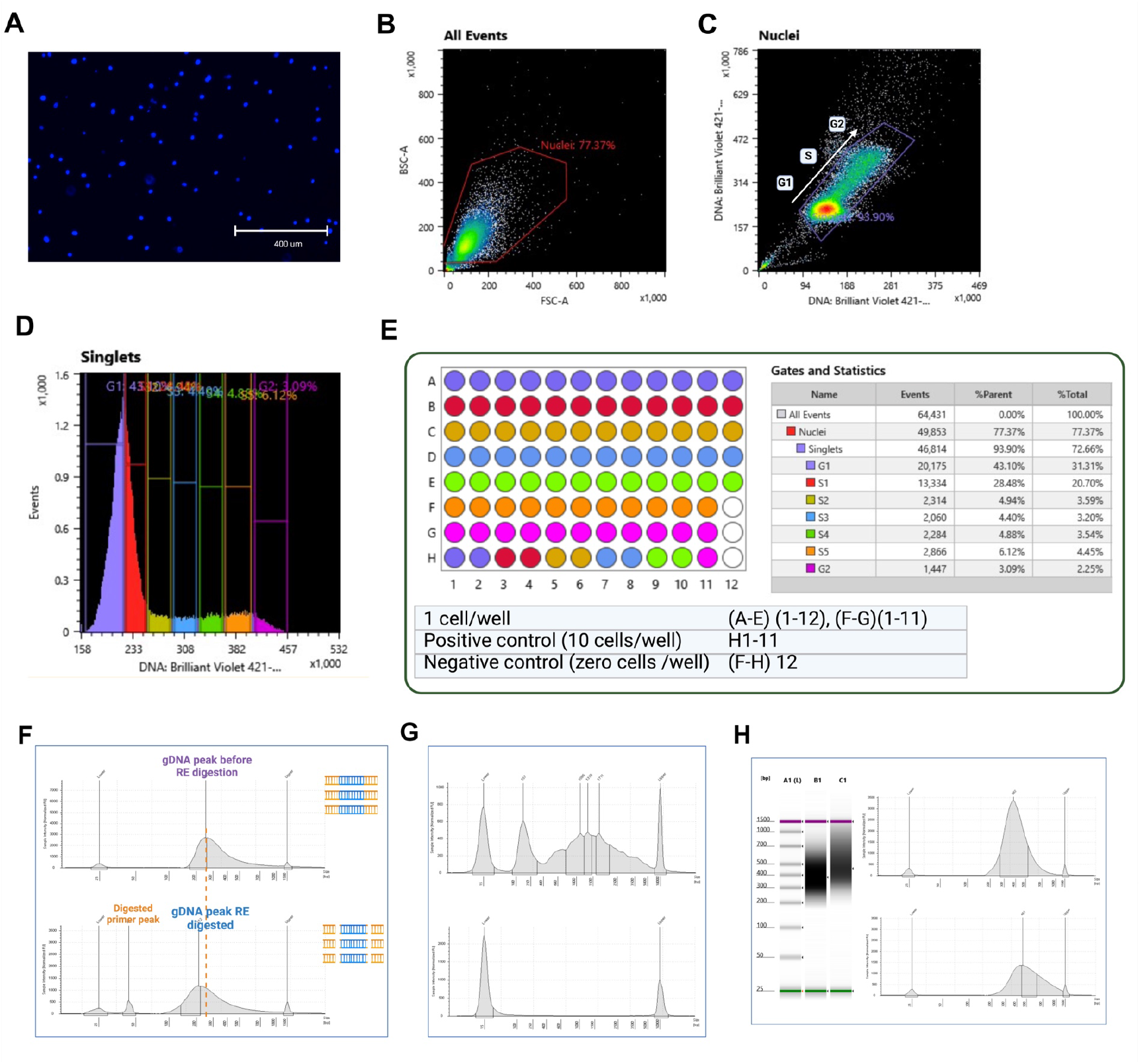
Sorting of Stained HepG2, for the Single Cell-Multiomics Protocol According to Cell Cycle. **(A)** HepG2 nuclei were stained with the DNA specific dye, Vybrant DyeCycle Violet. Sorting gate to capture **(B)** all events and **(C)** to capture stained nuclei in the G1, S and G2 phases of cell cycle. These were determined by the level staining of DNA dye on X axis and Y axis. **(D)** All events on Y axis with DNA specific staining on X axis. The sorting gates for G1, S, and G2 phase cells were set as shown. The S phase was divided into 5 equal gates from S1-S5. The identity of each gate is shown in the right panel in Figure E. (**E**) Design of 96-well plate sorted for the sc-multiomics protocol. <u>Left panel</u>-Design of a 96-well sorted plate. Single cells, and positive and negative control wells were included. <u>Right panel</u>-Gates sorted and colors assigned to each gate. Statistics of total events in each gate. **(F)** Amplified gDNA peaks before (top panel) and after RE digestion (bottom panel) detected using a Bioanalyzer, on a high sensitivity D1000 ScreenTape. Shift in the gDNA peak after RE digestion and appearance of the digested primer peak confirmed successful RE digestion. **(G)** Amplified cDNA peak detected using a Bioanalyzer, on a high sensitivity D5000 ScreenTape. The top panel demonstrates successful amplification of single cell cDNA with multiple peaks at 1.5-2 kbp. The lower panel shows a negative control well which had no nucleus. **(H)** Pooled gDNA and pooled cDNA wells were observed using a high sensitivity D1000 ScreenTape after library preparation and before sequencing. A1 is the ladder, B1 is the pooled gDNA samples and C1 is the pooled cDNA samples. <u>Top right pane</u>l-The pooled gDNA peak (B1) was observed at 382 bp. <u>Bottom right panel</u>-Pooled cDNA peak (C1) was observed at 461 bp.

When performing bulk Repli-seq, the S phase is sorted into ‘Early’ and ‘Late’ S-phase populations (Rivera-Mulia et al., 2022). With the single cell in-house approach, we gated the S phase population into 5 gates, S1 through S5 (**Fig. 2D**). This allowed us to track the gradual progression of genome replication through the S phase. For this study, we processed a total of 48 HepG2 nuclei across different cell cycle phases as described in the methods. The sorting design and cell cycle phase identity of each well is shown in **Fig. 2E**. We also processed positive and negative control wells.

Throughout the multiomics protocol, quality control (QC) was performed at regular intervals to ensure that no sample loss or contamination occurred. This also ensured successful amplification and processing of gDNA and cDNA at the different steps.

#### QC of amplified gDNA

We quantified the HepG2 gDNA per well using a Qubit. We obtained between 750-1000 ng of amplified gDNA per cell for 27 PCR amplification cycles. We also visualized amplified gDNA samples before and after RE digestion using high-sensitivity screen tapes on the Bioanalyzer. The amplified gDNA band was observed between 200-500 bp with a peak around 250 bp (**Fig. 4F, top panel**). After successful RE digestion, we observed a digested primer peak ∼45 bp along with a decrease in the gDNA fragments peak (from ∼250 to ∼205) (**Fig. 4F, bottom panel**). Absence of the primer peak at ∼45 bp indicated that RE digestion was not successful.

#### QC of amplified cDNA

We quantified the amplified cDNA using a Qubit. We found 22-24 amplification cycles were ideal for HepG2 cDNA amplification. We obtained 20-30 ng (22 cycles) and 75-100 ng (24 cycles) of cDNA per cell. We also visualized the amplified cDNA fragments on the Bioanalyzer. We observed a broad cDNA band between 500 to 4000 bp with multiple peaks around 1.5-2 kbp (**Fig. 4G, top panel**). The Smart-seq2 technique pulled down and amplified full length mRNAs of different lengths (Picelli et al., 2014). No peaks were observed in the cDNA negative control well (no nucleus) (**Fig. 4G, bottom panel**). Occasionally, we observed a peak around 150 bp, specific to the amplified primers. However, the bead purification steps after cDNA library preparation removed these primers.

We used a tagmentation-based library preparation kit for the amplified gDNA and cDNA, as described in the methods. We also employed uniquely barcoded oligos for gDNA and cDNA libraries, before pooling them together for sequencing.

### Copy number variation (CNV) and coverage in HepG2 single cell gDNA

We observed substantial genome coverage for all the sequenced sc-gDNA samples (**Fig. 3A**). Previously sequenced bulk HepG2 genome demonstrated aneuploidy in chromosomes 2, 6, 16, 17 and 20. Similar to the bulk genome, we observed chromosome specific CNV in the single cell genome plots (**Fig. 3A**). We also identified genes with CNV that were conserved across the HepG2 cells. These genes included TP53 binding protein *TP53BP2* (Huo et al., 2023), proto-oncogene *MYC (Chan et al*., *2004)*, Wnt regulator *CTNNB1* (He & Tang, 2020), PI3K pathway component *PIK3AP1* (AlGabbani, 2022), MAPK/ERK pathway kinase *BRAF* (Colombino et al., 2012), tumor suppressors *CDK2A, CDK2B and CDK2C* (Zhao et al., 2018), all of which are known to be mutated in the HepG2 cancer cells. A more complete listing of genes with CNV can be found in the attached supplemental file (**Supplemental file 1**).

**Figure 3:**
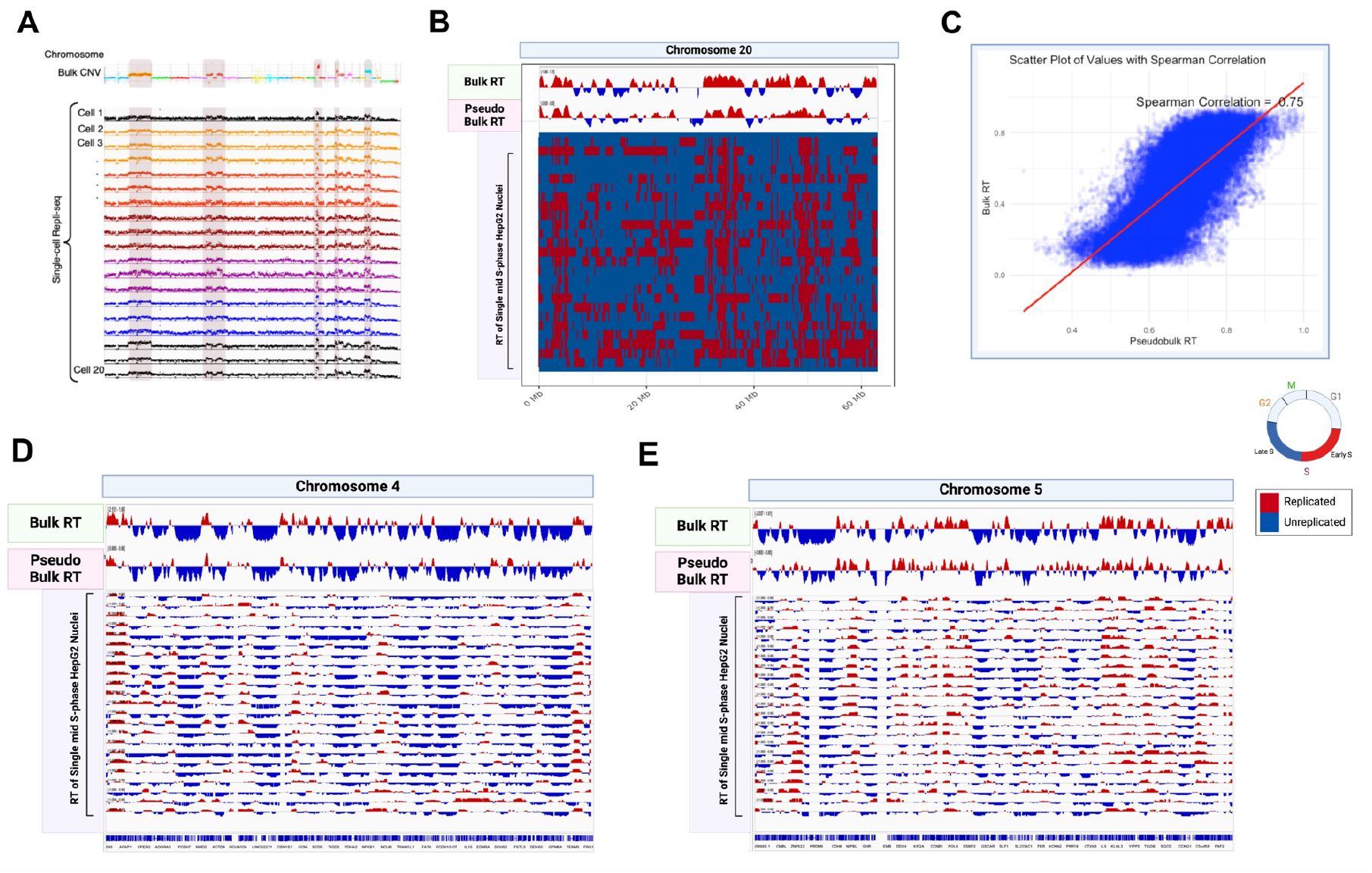
HepG2 Single Cell Replication Timing Profiles are Similar to HepG2 Bulk RT Profile. **(A)** HepG2 genome coverage for bulk (top panel) and single cell gDNA were generated from the multiomics protocol. CNVs in chromosomes 2, 6, 14, 16, and 20 are highlighted in the figure. **(B)** HepG2 Bulk, pseudo bulk of sc-RT profiles, and binarized sc-RT profiles for chromosome 20 are shown. Each row in the binarized plot is a S-phase single cell. (**C)** Spearman correlation of 0.75 was observed between genome-wide HepG2 bulk RT and pseudo bulk RT derived from sc-RT profiles. For **(D)** chromosome 4 and **(E)** chromosome 5, HepG2 bulk, pseudo bulk and non-binarized sc-RT profiles are provided. Replicated regions are highlighted in red and non-replicated regions are shown in blue.

### Analysis of single cell Replication Timing (sc-RT) in HepG2 Cells

We used established computational pipelines-Kronos sc-RT (Gnan et al., 2022) and the sc-Repliseq (Miura et al., 2020) for generating HepG2 sc-RT profiles as described in methods. The sc-RT profiles revealed conserved early or late replication domains across all cells, while also demonstrating smaller cell-to-cell variations. This could be observed in the binarized plot for Chromosome 20 (**Fig. 3B**) and in the non-binarized sc-RT plots for chromosome 4 and 5 (**Fig. 3D, 3E)**. Each row represents a sc-RT profile. Common patterns of replicated domains (red bins) were observed across all the cells. We also generated a HepG2 ‘pseudo bulk’ plot by combining the HepG2 sc-RT profiles of 17 single cells in the mid S-phase (described in the methods). The HepG2 pseudo bulk profile generated from 17 S phase single cells had a high correlation to the bulk reference RT profile, which was generated from tens of thousands of S-phase cells (**Fig. 3C**). Previous scRT protocols used ∼130 to 150 S-phase cells to deduce pseudo bulk RT (Gnan et al., 2022). Here, we demonstrated that with the gDNA processed using the in-house protocol, we could achieve significantly lower cell numbers. Since we sorted cells from the G1 phase, we were also able to observe the sc-RT domains switch from unreplicated to replicated, as cells transitioned from G1 to S phase (highlighted in **Supplemental Fig. 1C**).

### Gene expression of HepG2 nuclei

Using the sc-multiomics protocol, we processed and analyzed mRNA from the same HepG2 cells used for sc-RT analysis. We detected an average of 5586 unique genes per cell. We analyzed expression of cell cycle stage specific genes in the HepG2 cells sorted from G1, S and G2 phases. G2/M specific genes were significantly upregulated in the G2 cells compared to G1 and S phase cells (**Fig. 4A, top panel**). S phase specific genes were downregulated in G1 cells, upregulated in the S phase cells, and remained upregulated in the G2 population (**Fig. 4A, bottom panel**). This may have resulted from S-phase transcripts in the G2 phase, even in the absence of active transcription of S-phase genes.

**Figure 4:**
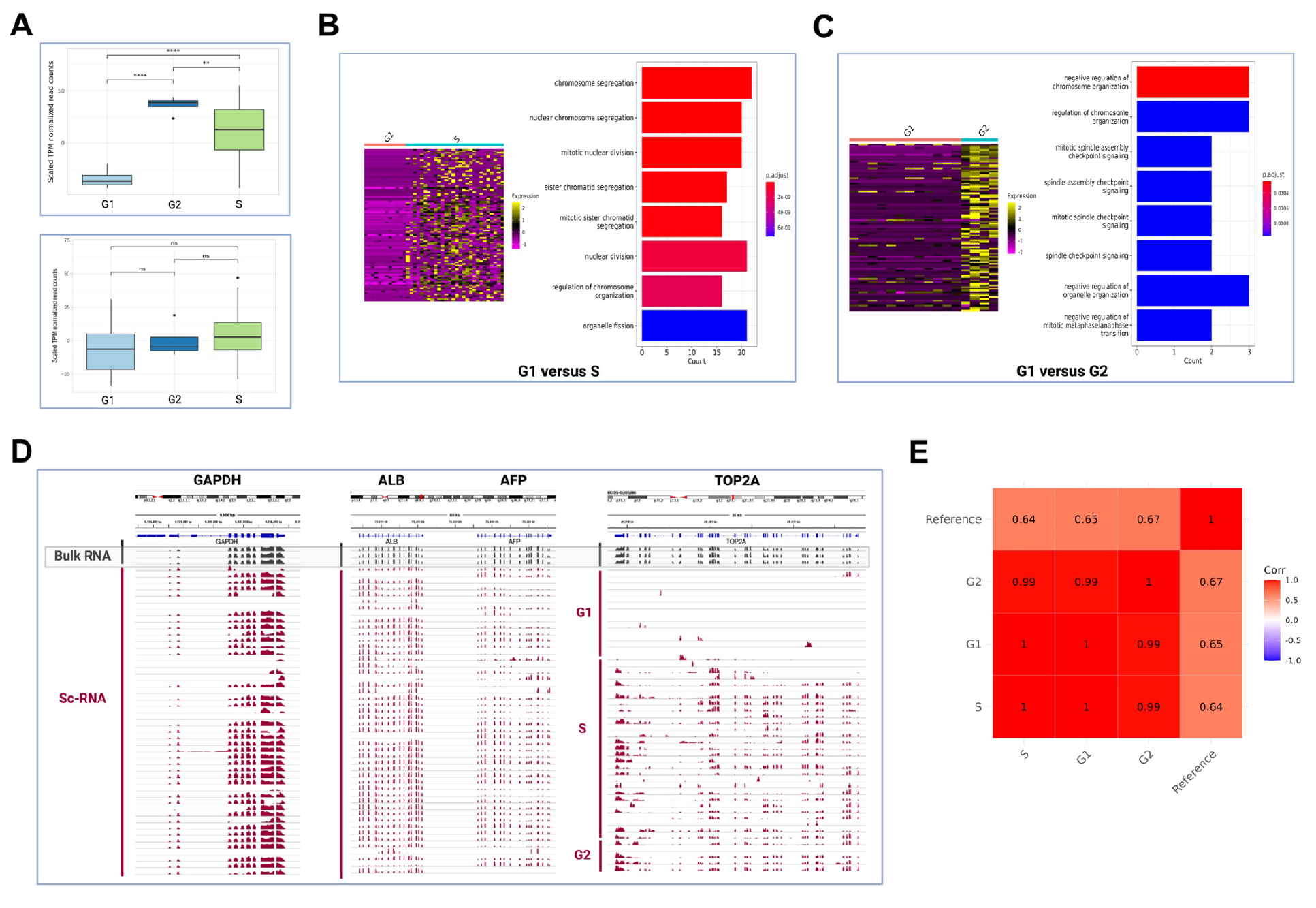
Gene Expression Analysis of scRNA Derived Using the Multiomics Protocol. **(A)** Level of expression of G2-phase specific genes (top panel) and S-phase specific genes (bottom panel) in populations of sorted G1, S, and G2 specific HepG2 cells. **(B)** Top differentially expressed genes (DEGs) between G1 and S cells and GO terms of top DEGS. **(C)** Top DEGs between G1 and G2 cells and GO terms of the top DEGs. **(D)** Comparison of gene expression between HepG2 bulk profiles and sc-RNA profiles. *GAPDH, ALB, AFP* were expressed across all populations. *TOP2A* was expressed in bulk, S and G2 cells but was not expressed in G1 cells. **(E)** Correlation between HepG2 scRNA from G1, S, G2 phases with RNAseq reference dataset (GSM923446).

We also generated heatmaps and analyzed gene ontology (GO) terms of the top 100 DEGs between stages. Top DEGs between G1 and S cells were associated with aspects of chromosome segregation, nuclear division, and chromosome organization, which are relevant as the cell transitioned from the G1 phase to DNA synthesis in the S phase (**Fig. 4B**). GO terms for G1 versus G2 phase were associated with chromosome organization and aspects of mitotic spindle checkpoint signaling (**Fig. 4C**). These analyses confirmed that the sc-RNA processing protocol allows us to capture a large number of unique genes, and conserved cell cycle stage-specific gene expression.

We observed high correlation between our HepG2 sc-RNA data and bulk HepG2 RNA seq data (**Fig. 4E**). We also compared housekeeping genes, hepatocyte specific genes, and cell cycle specific genes between the sc-RNA and bulk expression profiles (**Fig. 4D, Supplemental Fig. 2A)**. Expression of housekeeping genes like glyceraldehyde-3-phosphate dehydrogenase (*GAPDH*) and beta-actin gene (*ACTB*), as well as hepatocyte specific genes like albumin (*ALB*), alpha fetoprotein (*AFP*), fibrinogen (*FGB*) were conserved between the sc-RNA profiles and the bulk RNA profiles (**Fig. 4D, Supplemental Fig. 2A**). We also determined expression of cell cycle stage specific genes, cyclin-dependent kinase 1 (*CDK1*) and DNA topoisomerase II alpha (*TOP2A*). *CDK1* is highly expressed in G2/M phase but has very low or no expression in G1 phase (Liao et al., 2017). *TOP2A* is a proliferation marker that is expressed in the S/G2/M phases but has low expression or is absent in the G1 phase (Mjelle et al., 2015). Expression of *CDK1* and *TOP2A* were limited to single cells in the S and G2 phases only. No expression was observed in the G1 phase cells (**Fig. 4D, Supplemental Fig. 2A**).

### Correlation between RT and gene expression

Since we simultaneously extracted both gDNA and mRNA from the same cells, we were able to perform high resolution correlations between RT and gene expression (RT-RNA). Here, we performed RT-RNA correlations using (i) pseudo bulk RT and RNA for population trend; and (ii) sc-RT and sc-RNA for individual cell trends as described in the methods.

We observed a positive slope correlation between RT and RNA in the pseudo bulk analysis (**Fig. 5A**). Early replicating bins (>0.5) had significantly higher gene expression than late replicating bins (=<0.5) in the pseudo bulk RT-RNA analysis (**Fig. 5A, bottom panel**). We also generated the bulk HepG2 RT-RNA plot from reference RT and RNA datasets and observed the same positive slope trend (**Supplemental Fig. 2C**).

**Figure 5:**
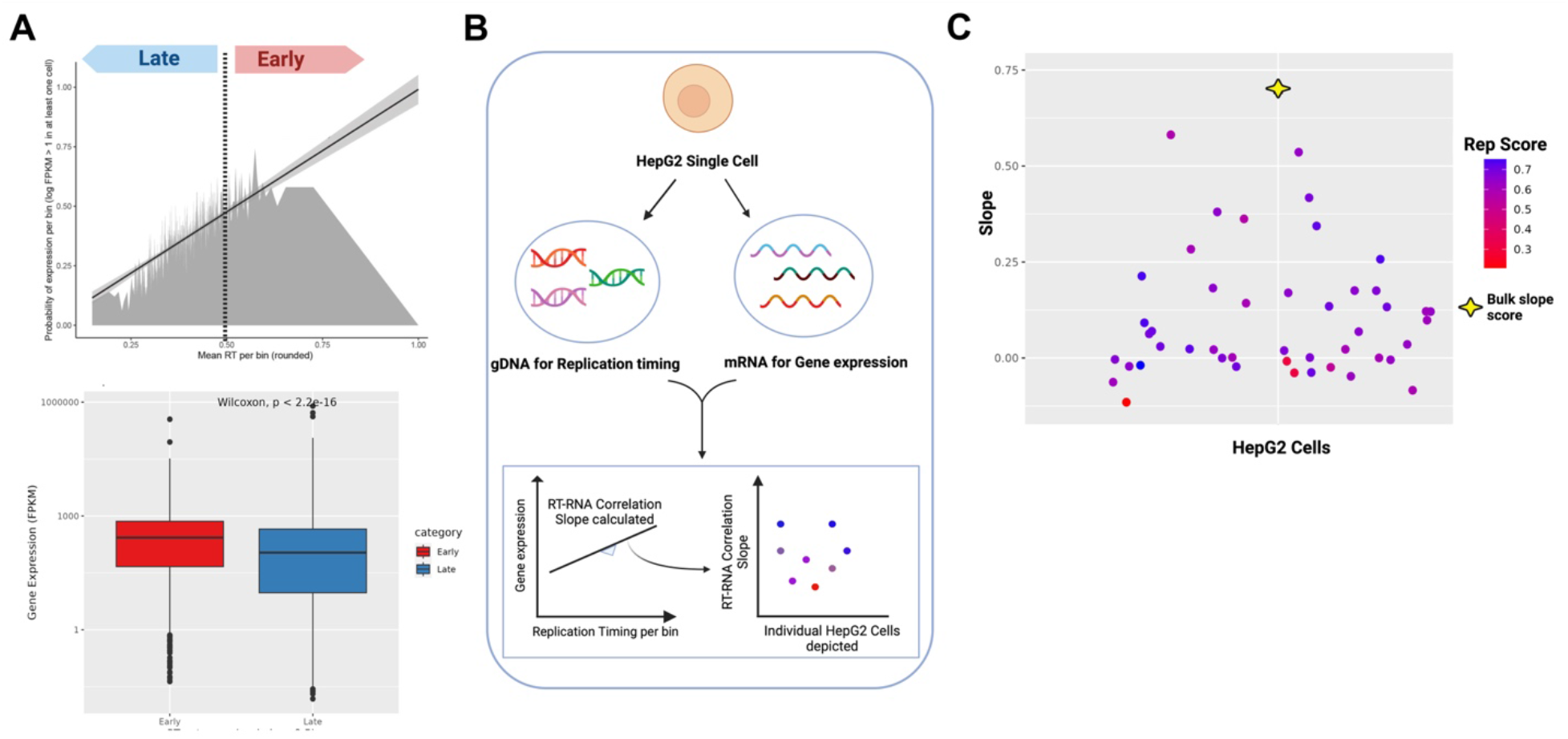
HepG2 Replication Timing (RT) and Gene Expression (RNA) Correlations at Pseudo Bulk and Individual Cell Level. **(A)** <u>Top panel</u> Pseudo bulk RT-RNA correlations plot with correlation slope. Bins are divided into early (>=0.5) and late replicating bins (<0.5). <u>Bottom panel</u> Significantly higher gene expression was observed in the early bins compared to late bins in the pseudo bulk RT-RNA analysis (Wilcoxon test, p < 2.2e-16). **(B)** Method used for deriving RT-RNA correlations within individual cells. Within each cell, the gene expression was plotted against RT and the slope was calculated. The correlation slopes of individual cells were then plotted for individual cells in figure **(C)**. The slope value of the bulk RT-RNA correlation is marked in the graph for comparison with the individual cells’ slopes.

Next, we analyzed RT-RNA correlations within individual cells and plotted the RT-RNA slope values for individual cells (**Fig. 5B**). We observed that the majority of cells had a positive RT-RNA correlation value, similar to the pseudo bulk analysis. We also plotted slope values against cell’s progress through S phase (Rep Score in **Supplemental Fig. 2B**) and observed similar trends. The level of RT-RNA correlation varied between cells in the same population. These cell-to-cell variations were lost with the pseudo bulk and bulk analyses. Using the sc-multiomics approach, we were able to capture single cell RT-RNA correlations at high resolution. Future studies can be focused on studying the different sub populations, based on RT-RNA scores, and analyzing how gene expression might vary between them.

## Discussion

We have developed an in-house single cell multiomics protocol to simultaneously analyze replication timing and gene expression, at the single cell level. This in-house protocol is much more affordable than the combined costs of commercially available kits for processing sc-gDNA and sc-RNA (∼15% of the kits’ costs for every 500 cells). Using this multiomics protocol, we extracted and amplified gDNA and mRNA from HepG2 nuclei. We analyzed genome coverage and observed CNVs at the chromosomal and gene levels. We also derived sc-RT profiles from gDNA and sc-gene expression information from mRNA. In cancer cells, changes in RT have been associated with aberrant gene expression and epigenetic changes (Donley & Thayer, 2013). Future studies can interrogate specific genes with CNV, and analyze their RT and gene expression, all within the same cells. This will allow us to draw more detailed correlations between the 3 parameters.

We observed that the pseudo bulk RT profile derived from the sc-RT profiles, had a high correlation to the HepG2 bulk RT profile. This allows us to drastically cut down on cell numbers compared to the bulk protocol. Here, we correlated RT with gene expression at the pseudo bulk level and the individual cell level. We were able to observe cell-to-cell variations which could not be observed in the bulk plots and underscores the ability of the sc-multiomics approach to capture cell specific differences.

While the protocol is compatible with cells or nuclei and can be used to process very small cell numbers, it is currently not high throughput. A future focus would be to customize the steps of the protocol to incorporate multiplexing and molecular barcoding to enhance high throughput. Another drawback is that we only gain insight into the mature RNA but not RNA without the poly A tail such as ribosomal RNA, microRNA, transfer RNA, and long non-coding RNA. gDNA and RNA separation techniques such as those used in scONE-seq (Yu et al., 2023) can be performed upstream of the in-house gDNA and mRNA amplification, to ensure that the whole transcriptome is captured.

The HepG2 multiomics dataset generated in this study can be used to generate gene regulatory networks to dissect molecular interactions at the single cell level. For example, we observed CNV in tumor suppressor genes (*TP53*), cell cycle regulator genes (*CDKN2A*), proto-oncogenes (*MYC*), genes involved in major pathways such as Wnt signaling (*CTNNB1*), PI3K/AKT pathway (*PTEN*), etc. all of which have been found to be mutated in HepG2 cancer cells as well as patient samples of hepatocellular carcinoma. This multiomics dataset can now be used to study CNVs of specific genes, and observe their gene expression and downstream targets, all within the same cell. This dataset can also be combined with single cell ATAC-seq and Hi-C datasets to gain insight into higher level chromosomal organization and relate it to RT and gene expression. In this study, all HepG2 nuclei were processed by hand using a multichannel pipette. However, the protocol can also be automated and performed using liquid handling robots such as EpMotion 5075, which will reduce manual processing time. An outline of the steps of the protocol are in **Supplemental Fig. 3**, which can be used as a reference for automation of the robot. More detailed steps are described in the methods.

## Methods

### Sorting of HepG2 nuclei based on cell cycle phase

We isolated and stained HepG2 nuclei with a DNA-specific dye based on a previously optimized protocol (Meyer-Nava et al., 2023). A total of 48 HepG2 nuclei were processed in this study from 2 different sorted plates (24 wells from each plate). The number of cells from each cell cycle phase gate are as follows-13 G1 phase cells, 6 S1 phase cells, 6 S2 phase cells, 5 S3 phase cells, 6 S4 phase cells, 6 S5 phase cells, 6 G2 phase cells.

### Single cell multiomics protocol for isolation and processing of gDNA and mRNA

The gDNA and cDNA were processed using the in-house multiomics protocol. Step-by-step description of the protocol is available at the protocol.io link in the data availability section. We used the Nextera XT DNA Library Prep Kit for tagmentation based library preparation of amplified gDNA and cDNA. To barcode each sample, we used unique oligos from the Nextera 96-index kit. All the gDNA and cDNA samples were then pooled on the same chip for sequencing. We used the NextSeq 2000 platform for sequencing-NextSeq P1 300 cycles, 150 PE. ∼2 million reads for gDNA and 1 million reads for cDNA were sufficient for deducing single cell RT and single cell gene expression profiles.

### Single cell replication timing analysis from HepG2 single cell gDNA

We used the Kronos scRT pipeline for generating the single cell RT binarized heatmaps. G1 and G2 sorted HepG2 cells were used as controls for normalizing gDNA reads from the S-phase HepG2 cells. We used bin sizes of 200kb for generating the scRT profiles on Kronos. We generated the scRT bed graphs showing non-binarized individual cell trends as well as the pseudo bulk plots using the sc-Repliseq pipeline (Miura et al., 2020). For the scRT bed graphs, we counted reads in 200kb sliding windows at 40kb intervals and then normalized reads against the HepG2 G1 control cells.

For the pseudo bulk plot, we considered sc-RT profiles of S2, S3 and S4 HepG2 nuclei (17 nuclei). Mid S-phase cells are ideal for deducing RT profile of a specific cell type. In mid S phase, half the genome is replicated and can be marked as ‘early’ while the non-replicated portion can be marked as ‘late’. Hence, for the pseudo bulk RT plot, we chose cells close to the mid S-phase, that is S2, S3 and S4 gated HepG2 cells. The reads from the S2-S4 cells were combined and processed as a single entry using the sc-Repliseq pipeline (Miura et al., 2020). Bulk HepG2 RT data was obtained from GSM923446. Copy number variation (CNV) was calculated as part of the Kronos scRT pipeline by loading the function ‘CNV’ which considers chromosomes of interest, chromosome size, genome wide bins. The CNV was adjusted per bin. Genes present in the bins that required adjustment are listed in the Supplemental file 1.

### Analysis of HepG2 gene expression

We used the Seurat v4 pipeline for the single cell RNA analysis and for generating the UMAP, heatmaps, and clusterProfiler for the GO analysis. The pseudo time trajectory analysis was done using Monocle3. Correlation between scRNA seq data generated in this study with reference scRNA database GEO-GSE90322 was performed.

### Correlation between Replication Timing and Gene Expression

Genes with FPKM>1 per stage were used for the correlation analyses. The RT of each gene was calculated from the pseudo bulk genome-wide RT plot for each stage. For RT bins values genome-wide, the probability of gene expression was calculated. For the individual cell correlation analysis, RT-RNA slopes were calculated within each cell, similar to the pseudo bulk analysis. The values of the slopes were plotted against a cell’s progression through S phase for individual cells.

## Data Availability Section

- All raw and processed sequencing data generated in this study have been submitted to the NCBI Gene Expression Omnibus (GEO; https://www.ncbi.nlm.nih.gov/geo/) under accession number GSE283896.
- All the code used for the analysis has been uploaded at GitHub link-https://github.com/AnalaShetty1/HepG2-scripts.
- The detailed steps for the single cell multiomics protocol are at protocols.io link-DOI: dx.doi.org/10.17504/protocols.io.36wgqdnwyvk5/v1

## Competing Interest Statement

The authors declare that they have no competing interests.

## Acknowledgments

We thank Dr. Juan Carlos Rivera-Mulia for early discussion and support on this project.

This work is supported by NIH grants R01 DK117286 (CJS), R01 DK117286-03S1 (CJS and WCL), R01 AI173804-01 (CJS and WCL).

Images were created using BioRender https://BioRender.com

## Notes

### Competing Interest Statement

The authors have declared no competing interest.

## References

AlGabbani, Q. (2022). Mutations in TP53 and PIK3CA genes in hepatocellular carcinoma patients are associated with chronic Schistosomiasis. Saudi Journal of Biological Sciences, 29(2), 848–853. 10.1016/j.sjbs.2021.10.022

Bartlett, D. A., Dileep, V., Baslan, T., & Gilbert, D. M. (2022). Mapping Replication Timing in Single Mammalian Cells. Current Protocols, 2(1), 1–27. 10.1002/cpz1.334

Briu, L.-M., Maric, C., & Cadoret, J.-C. (2021). Replication Stress, Genomic Instability, and Replication Timing: A Complex Relationship. International Journal of Molecular Sciences, 22(9). 10.3390/ijms22094764

Chan, K.-L., Guan, X.-Y., & Ng, I. O.-L. (2004). High-throughput tissue microarray analysis of c-myc activation in chronic liver diseases and hepatocellular carcinoma. Human Pathology, 35(11), 1324–1331. 10.1016/j.humpath.2004.06.012

Colombino, M., Sperlongano, P., Izzo, F., Tatangelo, F., Botti, G., Lombardi, A., Accardo, M., Tarantino, L., Sordelli, I., Agresti, M., Abbruzzese, A., Caraglia, M., & Palmieri, G. (2012). BRAF and PIK3CA genes are somatically mutated in hepatocellular carcinoma among patients from South Italy. Cell Death & Disease, 3(1), e259–e259. 10.1038/cddis.2011.136

Donley, N., & Thayer, M. J. (2013). DNA replication timing, genome stability and cancer. Late and/or delayed DNA replication timing is associated with increased genomic instability. In Seminars in Cancer Biology (Vol. 23, Issue 2, pp. 80–89). 10.1016/j.semcancer.2013.01.001

Gnan, S., Josephides, J. M., Wu, X., Spagnuolo, M., Saulebekova, D., Bohec, M., Dumont, M., Baudrin, L. G., Fachinetti, D., Baulande, S., & Chen, C. L. (2022). Kronos scRT: a uniform framework for single-cell replication timing analysis. Nature Communications, 13(1). 10.1038/s41467-022-30043-x

He, S., & Tang, S. (2020). WNT/β-catenin signaling in the development of liver cancers. Biomedicine & Pharmacotherapy, 132, 110851. 10.1016/j.biopha.2020.110851

Hiratani, I., & Gilbert, D. M. (2009). Replication timing as an epigenetic mark. Epigenetics, 4(2), 93–97. 10.4161/epi.4.2.7772

Hiratani, I., Ryba, T., Itoh, M., Yokochi, T., Schwaiger, M., Chang, C. W., Lyou, Y., Townes, T. M., Schübeler, D., & Gilbert, D. M. (2008). Global reorganization of replication domains during embryonic stem cell differentiation. PLoS Biology, 6(10), 2220–2236. 10.1371/journal.pbio.0060245

Huo, Y., Cao, K., Kou, B., Chai, M., Dou, S., Chen, D., Shi, Y., & Liu, X. (2023). TP53BP2: Roles in suppressing tumorigenesis and therapeutic opportunities. Genes & Diseases, 10(5), 1982– 1993. 10.1016/j.gendis.2022.08.014

Knowles, B. B., Howe, C. C., & Aden, D. P. (1980). Human hepatocellular carcinoma cell lines secrete the major plasma proteins and hepatitis B surface antigen. Science (New York, N.Y.), 209(4455), 497–499. 10.1126/science.6248960

Liao, H., Ji, F., & Ying, S. (2017). CDK1: beyond cell cycle regulation. Aging, 9(12), 2465–2466. 10.18632/aging.101348

Liu, L., De, S., & Michor, F. (2013). DNA replication timing and higher-order nuclear organization determine single-nucleotide substitution patterns in cancer genomes. Nature Communications, 4. 10.1038/ncomms2502

Macaulay, I. C., Haerty, W., Kumar, P., Li, Y. I., Hu, T. X., Teng, M. J., Goolam, M., Saurat, N., Coupland, P., Shirley, L. M., Smith, M., Van Der Aa, N., Banerjee, R., Ellis, P. D., Quail, M. A., Swerdlow, H. P., Zernicka-Goetz, M., Livesey, F. J., Ponting, C. P., & Voet, T. (2015). G&T-seq: Parallel sequencing of single-cell genomes and transcriptomes. Nature Methods, 12(6), 519–522. 10.1038/nmeth.3370

Macaulay, I. C., Teng, M. J., Haerty, W., Kumar, P., Ponting, C. P., & Voet, T. (2016). Separation and parallel sequencing of the genomes and transcriptomes of single cells using G&T-seq. Nature Protocols, 11(11), 2081–2103. 10.1038/nprot.2016.138

Marchal, C., Sasaki, T., Vera, D., Wilson, K., Sima, J., Rivera-Mulia, J. C., Trevilla-García, C., Nogues, C., Nafie, E., & Gilbert, D. M. (2018). Genome-wide analysis of replication timing by next-generation sequencing with E/L Repli-seq. Nature Protocols, 13(5), 819–839. 10.1038/nprot.2017.148

Meyer-Nava, S., Shetty, A. V, & Rivera-Mulia, J. C. (2023). Repli-seq Sample Preparation using Cell Sorting with Cell-Permeant Dyes. Current Protocols, 3(11), e945. 10.1002/cpz1.945

Miura, H., Takahashi, S., Shibata, T., Nagao, K., Obuse, C., Okumura, K., Ogata, M., Hiratani, I., & Takebayashi, S. ichiro. (2020). Mapping replication timing domains genome wide in single mammalian cells with single-cell DNA replication sequencing. Nature Protocols, 15(12), 4058–4100. 10.1038/s41596-020-0378-5

Mjelle, R., Hegre, S. A., Aas, P. A., Slupphaug, G., Drabløs, F., Sætrom, P., & Krokan, H. E. (2015). Cell cycle regulation of human DNA repair and chromatin remodeling genes. DNA Repair, 30, 53–67. 10.1016/j.dnarep.2015.03.007

Picelli, S., Faridani, O. R., Björklund, Å.K., Winberg, G., Sagasser, S., & Sandberg, R. (2014). Full-length RNA-seq from single cells using Smart-seq2. Nature Protocols, 9(1), 171–181. 10.1038/nprot.2014.006

Pope, B. D., Hiratani, I., & Gilbert, D. M. (2010). Domain-wide regulation of DNA replication timing during mammalian development. In Chromosome Research (Vol. 18, Issue 1, pp. 127–136). 10.1007/s10577-009-9100-8

Rivera-Mulia, J. C., Buckley, Q., Sasaki, T., Zimmerman, J., Didier, R. A., Nazor, K., Loring, J. F., Lian, Z., Weissman, S., Robins, A. J., Schulz, T. C., Menendez, L., Kulik, M. J., Dalton, S., Gabr, H., Kahveci, T., & Gilbert, D. M. (2015). Dynamic changes in replication timing and gene expression during lineage specification of human pluripotent stem cells. Genome Research, 25(8), 1091–1103. 10.1101/gr.187989.114

Rivera-Mulia, J. C., Desprat, R., Trevilla-Garcia, C., Cornacchia, D., Schwerer, H., Sasaki, T., Sima, J., Fells, T., Studer, L., Lemaitre, J. M., Gilbert, D. M., & Orr-Weaver, T. L. (2017). DNA replication timing alterations identify common markers between distinct progeroid diseases. Proceedings of the National Academy of Sciences of the United States of America, 114(51), E10972–E10980. 10.1073/pnas.1711613114

Rivera-Mulia, J. C., Trevilla-Garcia, C., & Martinez-Cifuentes, S. (2022). Optimized Repli-seq: improved DNA replication timing analysis by next-generation sequencing. Chromosome Research, 30(4), 401–414. 10.1007/s10577-022-09703-7

Štancl, P., Gršković, P., Držaić, S., Vičić, A., Karlić, R., & Korać, P. (2024). RNA-Sequencing Identification of Genes Supporting HepG2 as a Model Cell Line for Hepatocellular Carcinoma or Hepatocytes. Genes, 15(11), 1460. 10.3390/genes15111460

Vouzas, A. E., & Gilbert, D. M. (2023). Replication timing and transcriptional control: beyond cause and effect — part IV. In Current Opinion in Genetics and Development (Vol. 79). Elsevier Ltd. 10.1016/j.gde.2023.102031

Yu, L., Wang, X., Mu, Q., Sing, S., Tam, T., Shek, D., Loi, C., Chan, A. K. Y., Poon, W. S., Ng, H.-K., Chan, D. T. M., Wang, J., & Wu, A. R. (2023). scONE-seq: A single-cell multi-omics method enables simultaneous dissection of phenotype and genotype heterogeneity from frozen tumors. https://www.science.org

Zhao, Y., Chen, Y., Hu, Y., Wang, J., Xie, X., He, G., Chen, H., Shao, Q., Zeng, H., & Zhang, H. (2018). Genomic alterations across six hepatocellular carcinoma cell lines by panel-based sequencing. Translational Cancer Research, 7(2), 231–239. 10.21037/tcr.2018.02.14

