## Supplemental Figures for "Development of a Novel Single Cell Multiomics Approach for Simultaneous Analysis of Replication Timing and Gene Expression"

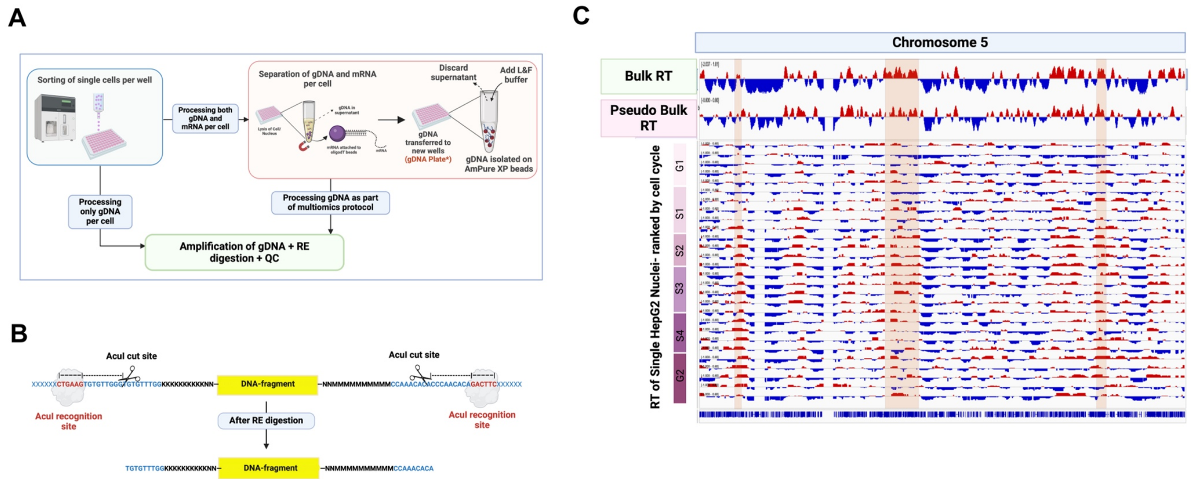


*Supplemental Figure 1:* **(A)** Overview of single cell genomic DNA (sc-gDNA) processing in-house protocol. The protocol can be used as part of the sc-multiomics protocol or as a stand-alone protocol to only process sc-gDNA. **(B)** Main strategy used to remove primer sequences after amplification of sc-gDNA and before sequencing of the sc-gDNA. **(C)** sc-RT plots of HepG2 cells ranked in order of cell cycle progression from G1 through G2 phases. The highlighted regions demonstrate regions where the switch from unreplicated (blue) to replicated (red) domains could be clearly observed with progress from G1 to S phase.

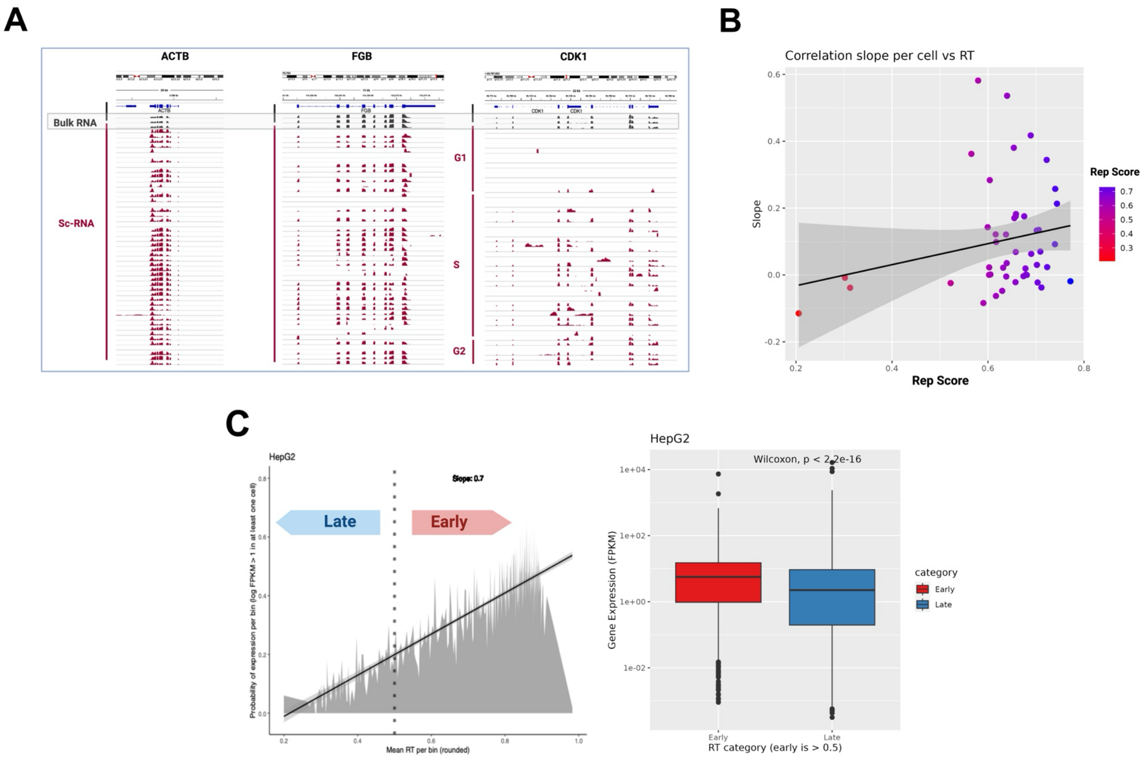


*Supplemental Figure 2:* **(A)** Comparison of gene expression between HepG2 bulk profiles and sc-RNA profiles. *ACTB* and *FGB* were expressed across all populations. *CDK1* was expressed in bulk, S and G2 cells but was not expressed in G1 cells. **(B)** RT-RNA correlation slope per cell plotted against Rep Score of cells. **(C)** HepG2 bulk RT-RNA correlation plotted from reference bulk RT (GSM923446) and bulk RNAseq (GSM923446) datasets. For the bulk RT-RNA correlation, the gene expression in early bins was significantly higher than in late bins (Wilcoxon test, p < 2.2e-16).

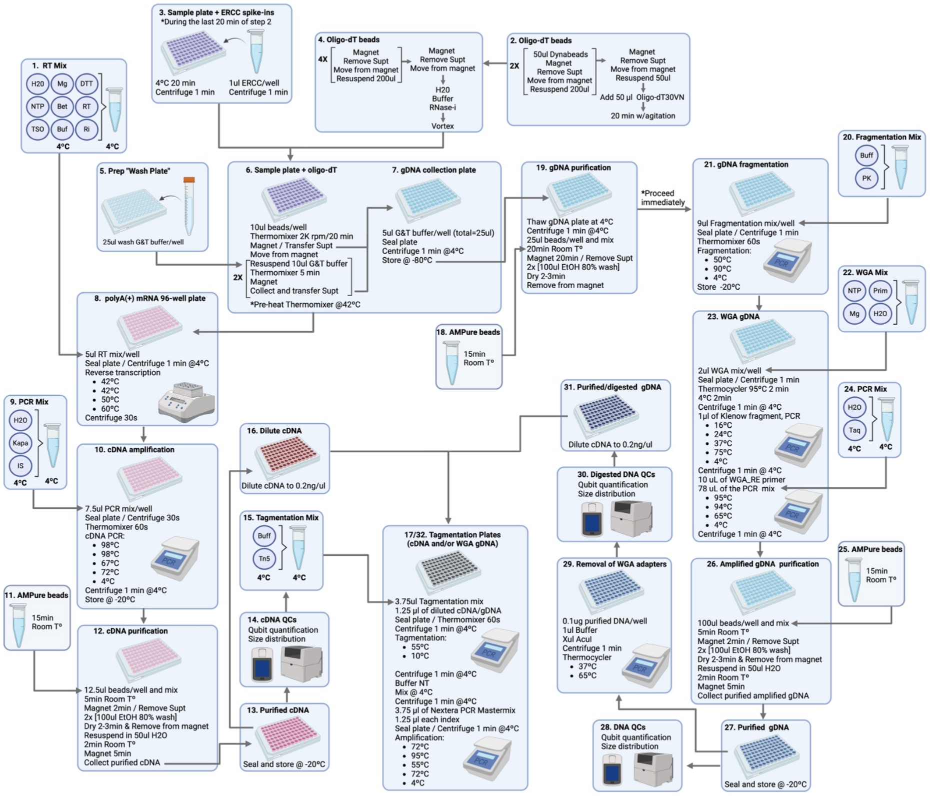


*Supplemental Figure 3: Step-by-step Protocol of the SC-Multiomics Approach. This figure can be used as a reference when designing automation of the sc-multiomics protocol on liquid handling robots such as the EpMotion 5075. Human intervention is required only when reagents need to be prepared fresh or when plates to be transferred to and from the centrifuge and thermocycler.*
